# Quartet protein reference materials and datasets for multi-platform assessment of label-free proteomics

**DOI:** 10.1101/2022.10.25.513670

**Authors:** Sha Tian, Dongdong Zhan, Ying Yu, Mingwei Liu, Yunzhi Wang, Lei Song, Zhaoyu Qin, Xianju Li, Yang Liu, Yao Li, Shuhui Ji, Yan Li, Lingling Li, Shanshan Wang, Proteomic Massive Analysis and Quality Control Consortium, Yuanting Zheng, Fuchu He, Jun Qin, Chen Ding

## Abstract

Quantitative proteomics is an indispensable tool in life science research. However, there is a lack of reference materials for evaluating the reproducibility of label-free liquid chromatography-tandem mass spectrometry (LC-MS/MS)-based measurements among different instruments and laboratories. We developed the Quartet as a proteome reference material with built-in truths, and distributed the same aliquots to 15 laboratories with nine conventional LC-MS/MS platforms across six cities in China. Relative abundance of over 12,000 proteins on 816 MS files were obtained and compared for reproducibility among the instruments and laboratories to ultimately generate proteomics benchmark datasets. There was a wide dynamic range of proteomes spanning ~7 orders of magnitude (10^1^–10^8^ copies/cell), and the injection order had marked effects on quantitative instead of qualitative. Overall, the Quartet offers valuable standard materials and data resources for improving the quality control of proteomic analyses as well as the reproducibility and reliability of research findings.

## Introduction

The launch of the Human Phenome Project^1–4^ and the Human Proteome Project has become the consensus of the scientific community in the post-genomic era and brought the field of proteomics to a prominent position in the life sciences owing to its implications in precision medicine^5–9^. Continuous improvements in next-generation proteomics, such as in instrumentation, sample preparation, and computational analysis, have facilitated the generation of large amounts of data, including protein profiles, post-translational modifications, and protein-protein interactions^10–13^. Similar to other omics technologies (e.g., genomics and transcriptomics), low reproducibility across large-scale experiments remains one of the most critical challenges in proteomics^14, 15^. This poor reproducibility is mainly attributable to the large variability introduced by heterogeneity in experimental design, sample processing methods, environmental and operating conditions, mass spectrometers, data search algorithms, statistical analysis methods, and other conditions across studies and laboratories^16^. Moreover, there has been no comprehensive comparison of proteomics datasets obtained using different liquid chromatography-tandem mass spectrometry (LC-MS/MS) systems and conditions, posing a challenge for comparative proteomics and subsequent biological interpretation. Thus, there is an urgent need to develop standard reference materials, establish standard operating procedures (SOP), and generate standard reference datasets for comprehensive quality control (QC) analysis in proteomics.

The US Food and Drug Administration (FDA)-led Microarray and Sequencing Quality Control (MAQC/SEQC) consortium conducted three phases of projects to assess the reliability and reproducibility of genomics technologies, including microarrays, genome-wide association studies, and next-generation sequencing^17–19^. At the end of 2017, the MAQC consortium announced the formation of the MAQC Society, an international society dedicated to the quality control and analysis of massive data generated from high-throughput technologies with the goal of enhancing reproducibility^20^. In addition, the Microbiome Quality Control (MBQC) project was established to evaluate and, ultimately, standardize measurement methods for the human microbiome and includes protocols for handling human microbiome samples and computational pipelines for microbial data processing^21^.

Similarly, tremendous effort has been made to acquire high-quality and reproducible data in the field of proteomics. Paulovich et al.^22^ put forward an SOP for the large-scale production of yeast standards and offered aliquots to the proteomics community through the National Institute of Standards and Technology, where the yeast proteome is under development as a certified reference material to meet the long-term needs of the community. In cooperation with the Human Proteome Organization Test Sample Working Group, Bell et al.^23^ performed a test sample study to identify errors—such as incompleteness of peptide sampling—leading to irreproducibility in LC-MS/MS-based proteomics. The authors distributed an equimolar test sample comprising 20 highly purified recombinant human proteins to 27 laboratories. The centralized analysis determined missed identification (false negatives), environmental contamination, database matching, and curation of protein identification as sources of problems. To determine the inter- and intra-laboratory reproducibility and performance of sequential window acquisition of all theoretical fragment ion spectra (SWATH)-MS for large-scale quantitative proteomics, Collins et al.^24^ created a benchmarking sample set comprising 30 stable isotope-labeled standard peptides diluted into a complex background, and then distributed the aliquots to 11 laboratories worldwide to assess the reproducibility of quantitative proteomics data. Using SWATH-MS data acquisition, the reproducibility, limit of detection, and linear dynamic range of quantitative characteristics were highly comparable across data from all sites.

Despite these efforts, there has been no baseline investigation similar to that of the MAQC/SEQC and MBQC for the proteomics community to systematically evaluate data reproducibility across laboratories and identify potential measurement variability in each step of the general proteome workflow. Such comparative analyses have been hindered by several factors, including (i) lack of a uniform standard reference material for proteomics analyses, (ii) variation among laboratories in the methods used for sample preparation, (iii) variations in data quality due to the use of different data acquisition methods, and (iv) lack of a consistent evaluation system for subsequent analysis of the generated proteomics data^25–27^. These factors have substantially impeded the ability to translate proteome technology into actionable clinical practice. Therefore, launching a proteome QC project to establish uniform proteome standard reference materials, formulate uniform proteome SOPs, and generate uniform proteome benchmark datasets is imperative and will facilitate assessment of variations in proteome analysis processes, help determine the advantages and limitations of various MS instruments and bioinformatics strategies, enhance consensus on data analysis methods, and ultimately improve the robustness of results.

To address these issues, we initiated the proteome QC project as part of MAQC phase IV (MAQC-IV). In this study, we developed the Chinese Quartet (hereafter referred to as “the Quartet”) as a proteome standard comprising four standard samples derived from four members of the same family. For establishing the Quartet, intrinsic differences between samples were defined as “built-in truths,” which enabled evaluation of the qualitative and quantitative characteristics of LC-MS/MS-derived data. We compared the proteins identified in all measurements of each sample of the Quartet and identified differentially expressed proteins (DEPs) across experiments with three technical replicates. The performance of different platforms was quantitatively evaluated using the signal-to-noise ratio (SNR) to determine variations and differences among samples. We further evaluated batch effects due to differences in the injection order of LC-MS/MS and performed absolute quantification of the Quartet standards using C^13^ stable isotope-labeled concatenated peptides (QconCAT). As a result, we provide a standard reference material for evaluating the reproducibility and reliability of different MS-based proteomics strategies. The variations introduced by different factors, including MS principles, instrument specs, and temporal and spatial issues, provide a valuable resource for the development and optimization of proteomics SOPs that are applicable to both basic and clinical studies^28^.

## Results

### Establishment of the Quartet

Lymphoblastoid cells provided by a Chinese family from the Fudan University (FDU) Taizhou cohort^29–31^, including the father (F7), mother (M8), and a pair of twin daughters (D5 and D6), were used to establish immortalized cell lines. Protein extraction from cells and peptide digestion were uniformly performed in the National Center for Protein Sciences (NCPSB, Beijing, China) laboratory. We distributed these aliquots to 15 laboratories throughout six cities in China (Fig. 1a). Nine types of conventional LC-MS/MS platforms, including Orbitrap-based systems (e.g., Orbitrap Lumos and Q-Exactive) or time-of-flight (TOF)-based systems (TripleTOF 6600 and timsTOF pro), were incorporated in the proteomics platform evaluation. All raw LC-MS/MS files were uploaded into Firmiana^32^, and quantitative proteome matrix results were obtained following the standard computational workflow for proteomics data. The assessment for multi-character datasets was performed by the bioinformatics team at the FDU laboratory. The advantage of this strategy was that any factors influencing the identification and quantification of (differentially expressed) proteins in proteomics data were restricted to the different sample types, sites, and instruments, excluding any variations in sample preparation and downstream bioinformatics methods.

**Fig. 1.**
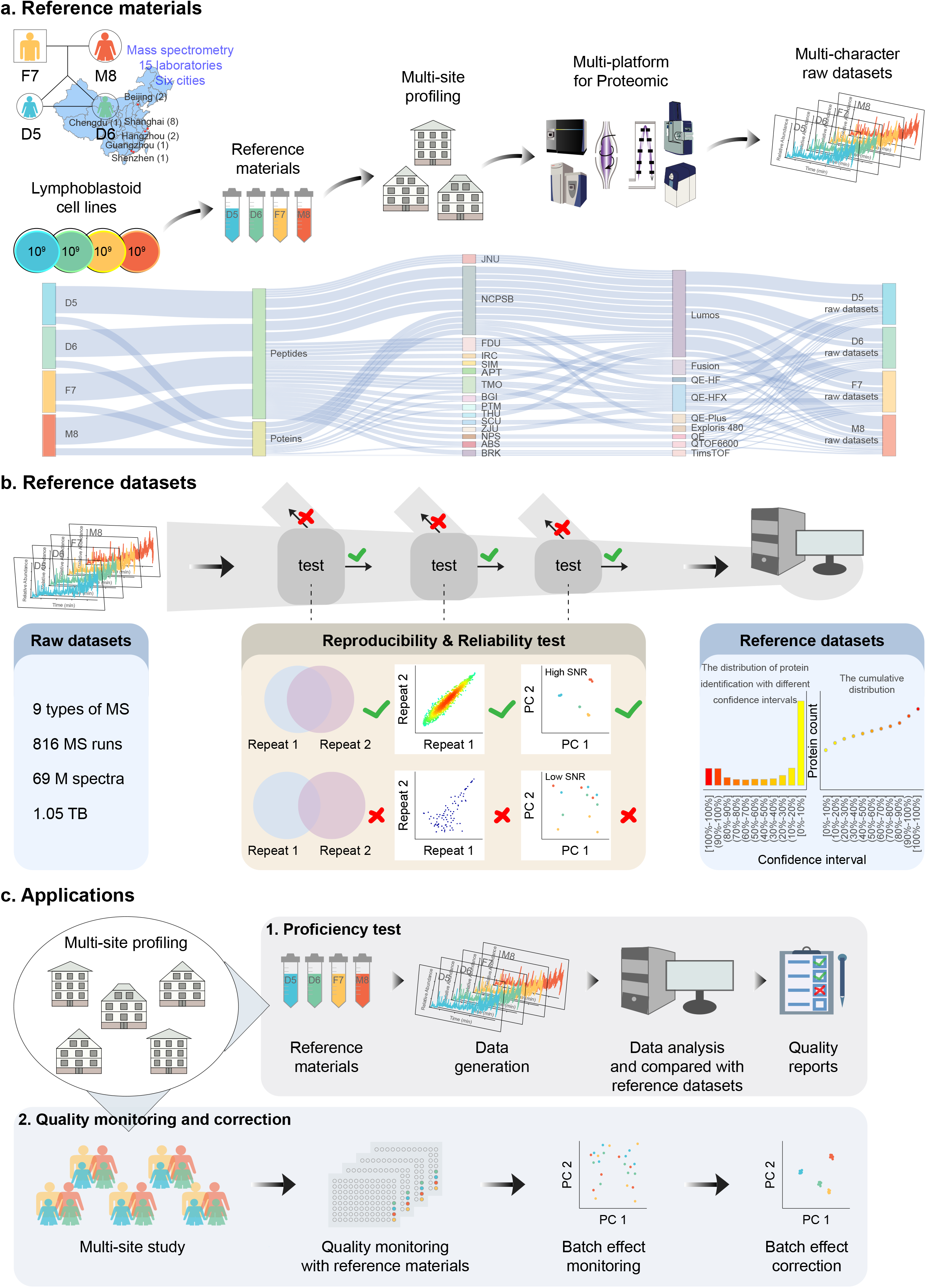
Study design and implementation. **a**, Lymphoblastoid cells provided by a family, consisting of a father (F7, yellow), mother (M8, red), and a pair of twin daughters (D5 and D6, apple green and azure), were used to prepare the proteome reference materials (i.e., the Quartet) in both protein and peptide forms. The aliquots were sent to 15 laboratories, and proteomics analyses by data-dependent acquisition (DDA) was carried out using Orbitrap- and TOF-based systems. All LC-MS/MS files were aggregated at Fudan University for analysis. **b**, A total of 816 MS files were used to qualitatively and quantitatively evaluate the reproducibility and reliability of the proteome. Furthermore, reference protein sets with confidence intervals were determined. **c**, Application scenarios of the reference materials.

All measurements resulted in a global dataset containing 816 MS files generated from the 15 laboratories (Supplementary Table 1), providing a rich resource for performing both qualitative and quantitative reproducibility and reliability assessments while promoting the clinical translation and application of proteomics. Among the files, 792 MS files were used for the following purposes: cross-platform assessment with a single-shot proteomics strategy, which acquired a total of 288 LC-MS/MS runs (4 samples × 3 replicates × 24 experiments) using the Quartet standards at the peptide level (Supplementary Fig. 1a); deep coverage with a sample fractionation strategy, which produced a total of 384 LC-MS/MS runs using the Quartet standards at the protein level (Supplementary Fig. 1b); and longitudinal proteomic monitoring for 15 months, which was used for sample stability evaluation, generating 120 MS files using the Quartet standards at both peptide and protein levels (Supplementary Fig. 1c).

The reference material (i.e., the Quartet) allowed us to perform daily QC tests. Using the entire dataset, we further identified reference protein sets with confidence intervals to provide important benchmarks for the application of LC-MS/MS technologies to clinical assays. The confidence intervals were determined according to the frequency of protein occurrence in all experiments. These reference datasets can offer a comparable baseline for guiding users to carry out proficiency tests, promoting parameter optimization and method development. The Quartet can also be used to assess the reproducibility and performance of large-scale quantitative proteomics among different instruments and laboratories. The built-in individual differences can facilitate evaluation of the reproducibility of differential proteome platforms (Fig. 1b, c).

### Characteristics of the Quartet

To delineate the proteomic portraits of the reference materials, we performed 110 MS runs on each sample, including 102 single-shot and 8 deep-coverage MS runs. Over 10,000 proteins with at least one unique and strict peptide were detected in all measurements of each Quartet sample. Up to 12,068 proteins were identified in the Quartet using state-of-the-art high-resolution MS, indicating that MS technology is capable of a coverage of 10,000 proteins per sample (Fig. 2a, b). The deep-coverage strategy with multiple fractions (≥ 6) detected ~4,000 more proteins than did the singleshot strategy. Gene ontology enrichment analysis (http://geneontology.org) for cellular components of the Quartet proteome mainly resulted in nine terms, covering the nucleus, cytosol, plasma membrane, cytoskeleton, mitochondrion, extracellular space, Golgi apparatus, endoplasmic reticulum, and lysosome, demonstrating the complexity and diversity of the reference materials (Fig. 2c). In the 24 experiments (with three technical replicates) using the single-shot strategy, the average number of proteins identified in each sample varied from 1,500 to 5,000 depending on the different mass spectrometers used in the 15 test sites (Fig. 2d). For example, ~2,000 proteins were detected with the Q Exactive and TripleTOF 6600 instruments, whereas over 4,500 proteins were identified with the Exploris 480 and timsTOF Pro instruments. Notably, Site 11 identified less than 2,000 proteins with Q Exactive Plus, which was highly inconsistent with the empirical value, suggesting that the state of the LC-MS/MS system should be inspected or that an inferior proteomics workflow was carried out. In the eight fractionation experiments, with 6, 12, 18, or 30 fractions, over 4,500 proteins were identified in each sample, ranging from 4,745 proteins identified with the TripleTOF 6600 system using six fractions to 8,441 proteins identified with the Fusion Lumos system using 30 fractions (Fig. 2e).

**Fig. 2.**
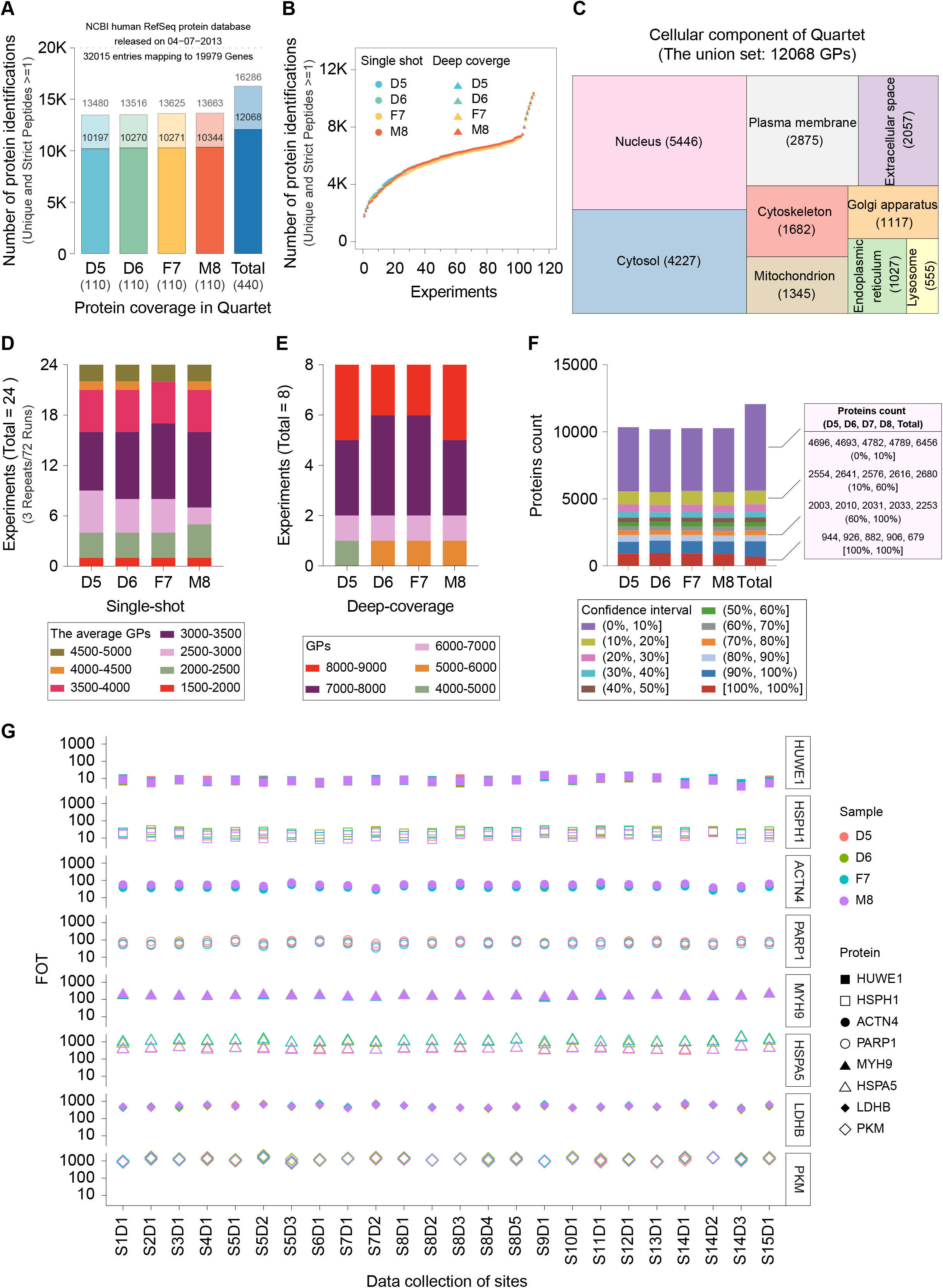
Characteristics of the Quartet. **a**, Proteins identified from the Quartet samples. **b**, Cumulative number of proteins identified as a function of experiment numbers. **c**, Gene ontology analysis of cellular component terms (GO-CC). **d**, Average number of proteins identified from different experiments using the single-shot strategy. **e**, Number of proteins identified from different experiments using a deep-coverage strategy. **f**, Number of proteins identified distributed over 11 confidence intervals. The stack column labeled “Total” indicates the global distribution of all proteins identified from the Quartet samples. Right panel: detailed protein identification information within rough confidence intervals: 0–10%, 10–60%, 60–100%, and 100–100%. **g**, Proteins with high quantitative stability distributed into different quantitative orders of magnitude.

We divided the proteins identified from 110 MS runs of each Quartet sample into 11 groups with different confidence intervals according to their occurrence frequency in all experiments (observation times/total measurement times). For each sample, proteins with a confidence interval of 0–10%, 10–60%, 60–100%, and 100–100% accounted for approximately 46%, 25%, 20%, and 9% of the total, respectively. In the whole proteome (12,068 proteins) of the Quartet, 679 proteins (6%) with ultra-high confidence were identified in all 440 MS runs (Fig. 2f), including some proteins with high quantitative stability that were distributed into different quantitative levels (Fig. 2g), such as HUWE1, HSPH1, ACTN4, PARP1, MYH9, HSPA5, LDHB, and PKM, which have potential value as internal “anchor” proteins for quantification. During quantitative analysis of a complex clinical proteome, the calibration of protein profiles in the experiment according to these anchor proteins may improve the reliability of the detected protein abundance.

### Differential proteome as a reliability indicator

The Quartet was designed to include built-in truths among the samples of the four family members to discover quantitative differences across proteome platforms. In the whole Quartet proteome (12,068 proteins), 74% (8,934) of the proteins were common to the D5, D6, F7, and M8 samples. In contrast, each sample contained ~4% specific proteins, ranging from 394 proteins in D6 to 421 proteins in M8 (Fig. 3a). Differential proteomics analysis was then performed for the 24 experiments (with three technical replicates) using the single-shot strategy. Using a fold-change threshold of > 2 observed at least once in pairwise comparisons of the four Quartet samples within a single experiment, a total of 1,662 DEPs were identified across all experiments, demonstrating the sample-specific characteristics that can be used for constructing built-in truths of reference materials (Fig. 3b). Even though samples D5 and D6 had identical genomes, they exhibited differential gene expression patterns at the proteome level, reflecting the complexity and variations in the transcription and translation processes among individuals.

**Fig. 3.**
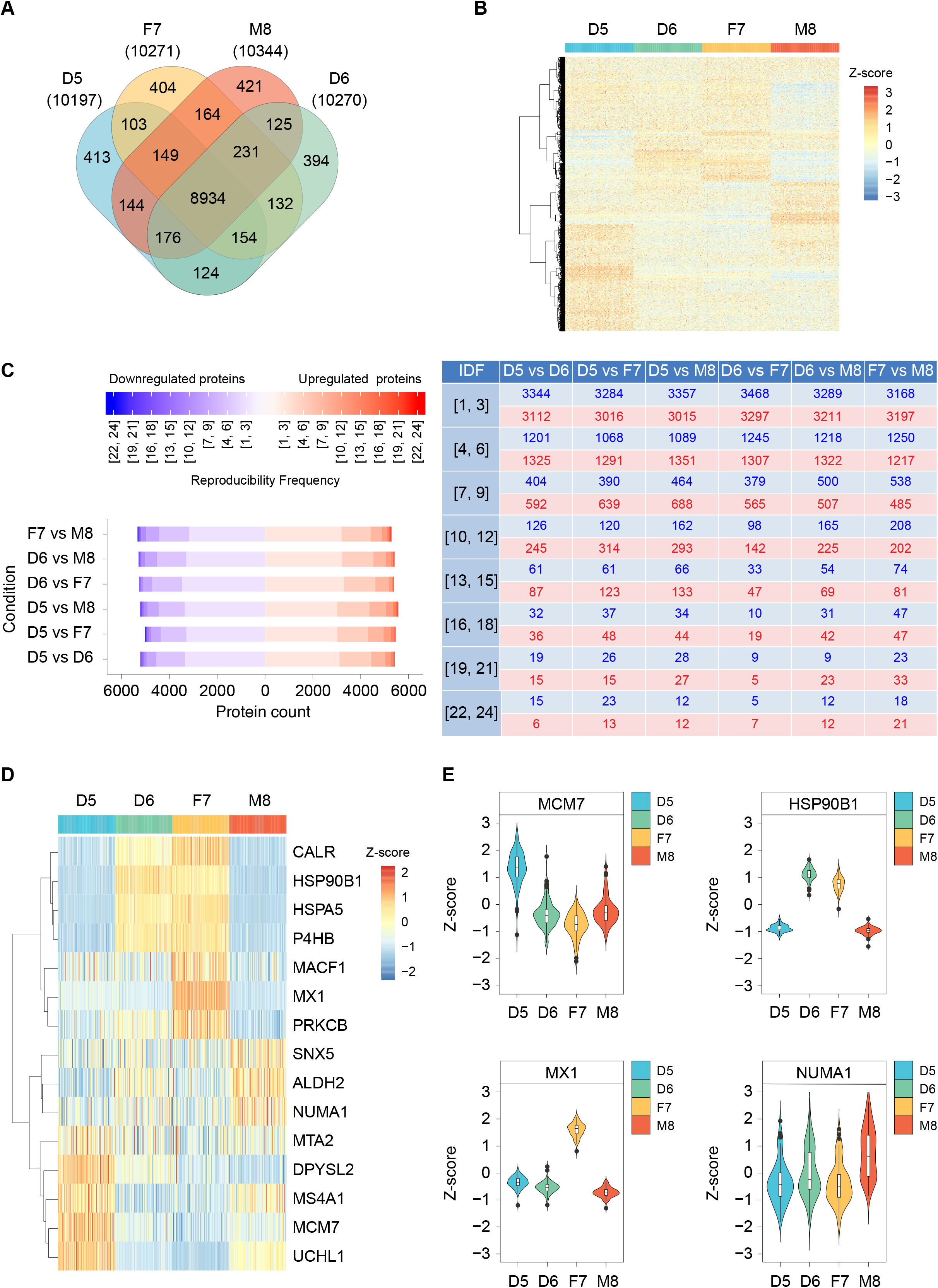
Differential proteome as a reliability indicator. **a**, Venn diagram of proteins identified in the Quartet samples. **b**, Heatmap of differentially expressed proteins. Cluster analysis was performed using Z-score-transformed protein intensities for the proteins with expression level fold change > 2. Red indicates a high expression level and blue indicates a low expression level. **c**, Occurrence frequency of differentially expressed proteins (DEPs), according to the pairwise comparisons between samples of the Quartet, over 24 experiments with three technical replicates each using a single-shot strategy. Left panel: number of DEPs in different partition intervals for each pairwise comparison. Red and blue indicate upregulation and downregulation directions of DEPs within a comparison, respectively, such as D5 versus D6, with D6 as the reference group. Right panel: numeric table corresponding to the distribution map in the left panel. IDF: identification frequency. **d**, Heatmap of 15 DEPs with a consistent upregulation/downregulation trend across all 24 experiments. **e**, Proteins with highly sample-specific expression in the Quartet.

Furthermore, to determine the high-confidence DEPs that can serve as “indicators” to evaluate the quantitative reliabilities of different MS platforms, we counted the occurrence frequency of DEPs in the 24 experiments by pairwise comparisons between samples within each experiment, which was considered to reflect the reproducibility frequency of DEPs. We divided the reproducibility frequency of DEPs into eight intervals as shown in Fig. 3c. The results indicated 5,418 upregulated proteins and 5,202 downregulated proteins (DEP pool) in D5 compared with D6 in at least one experiment (Supplementary Table 2 and Table 3), of which 94.0%, 5.5%, and 0.5% exhibited a low (1-9), medium (10-18), and high (19-24) reproducibility frequency, respectively. After applying the same analysis for other pairwise comparisons within the family, it became apparent that the main source of variation causing low reproducibility is likely different mass spectrometer models, different test sites, and random MS/MS sampling by data-dependent acquisition. DEPs with medium (10-18) and high (19-24) reproducibility frequencies showed high robustness, which generally reflected real differences between the samples in the Quartet. These proteins can be used as a benchmark dataset to evaluate the quantitative ability of comparative proteomics on different platforms. Additionally, we found 15 DEPs with consistent upregulation/downregulation trends in all 24 experiments (Fig. 3d); for example, MCM7, HSP90B1, MX1, and NUMA1 exhibited significantly high expression levels in D5, D6, F7, and M8, respectively (Fig. 3e). Overall, the detection rates and differential expression patterns of these proteins can serve as a baseline for evaluating the performance of different LC-MS/MS platforms in proteome detection.

### Variation among MS instruments

Performance comparison of different instruments can provide guidance for optimizing experimental strategies for users while highlighting technological upgrades for manufacturers. We thus analyzed 108 MS files (9 LC-MS/MS × 4 samples × 3 repeats) produced by nine conventional instruments by detecting the Quartet standards in terms of qualitative and quantitative aspects. These instruments were mainly categorized into Orbitrap-type (Fusion series, Fusion and Fusion Lumos; QE series, Q-Exactive, Q-Exactive Plus, Q-Exactive HF, Q-Exactive HF-X, and Exploris 480 with ion mobility) and TOF-type (TripleTOF 6600 and timsTOF Pro with ion mobility) mass spectrometers. We performed four comparisons among the following instruments: (1) two Fusion series instruments, (2) five QE series instruments, (3) two TOF-type instruments, and (4) two instruments equipped with ion mobility. Taking sample D5 as an example, Fusion Lumos identified 4,425 proteins, which was 200 proteins more than those identified with Fusion. In the QE series, the five instruments possess upgraded scanning speeds and other configurations introduced with each generation, and the number of proteins identified also gradually increased in parallel with the model upgrade. For example, 2,777 proteins were detected using the first generation of QE (Q-Exactive) while 5,248 proteins were detected with the latest generation (Exploris 480). For the TOF series, TripleTOF 6600 identified 2,735 proteins, whereas timsTOF Pro identified 5,194 proteins. Thus, timsTOF Pro and Exploris 480, which are both equipped with ion mobility capability, resulted in comparable protein identification (5,194 and 5,248, respectively), indicating the advantages of ion mobility in proteome screening (Fig. 4a).

**Fig. 4.**
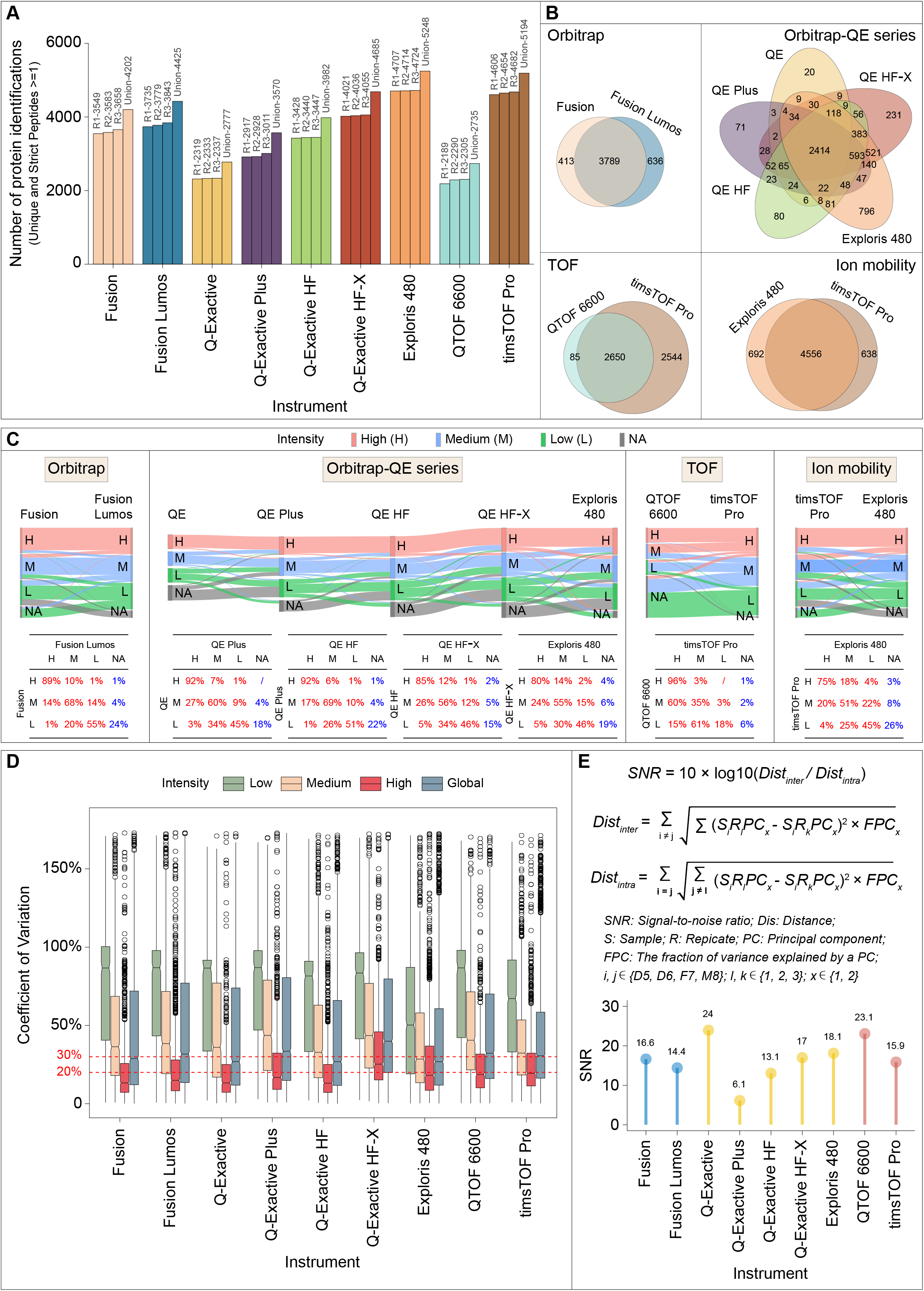
Variation among nine conventional instruments. **a**, Number of proteins identified in sample D5 (one of the twin daughters) from the Quartet using nine conventional instruments. **b**, Venn diagram of proteins quantified from the same series of MS instruments, corresponding to Orbitrap Fusion series, Orbitrap QE series, TOF series, and ion mobility series. **c**, Sankey plot (top panel) depicting the flow direction of reproducibility for proteins within different (low, medium, high) intensity groups along with upgraded instruments. The corresponding and detailed percentages are summarized in the bottom table. This assessment involved the same four series of MS instruments as described for (**b**). For a specific experiment, low intensity refers to the < 33.33th percentile, medium intensity refers to the 33.33–66.7th percentile, and high intensity refers to the > 66.67th percentile. **d**, Coefficients of variation (CVs) for proteins within different intensity groups and the global group for the nine conventional instruments (also see Fig. S2b). **e**, Definition of signal-to-noise ratio (SNR, top panel) and the SNR distribution of the nine conventional instruments (bottom panel).

Furthermore, we delineated the protein identification reproducibility of the Fusion, QE, and TOF series and mass spectrometers with embedded ion mobility. As shown in the Venn diagram in Fig. 4b, 90% (3,789) of the proteins identified by Fusion were also identified by Fusion Lumos; Exploris 480, as the most advanced instrument in the QE series, covered more than 87% (4,452) of the proteins identified by the previous four generations of instruments; timsTOF Pro identified 97% of the proteins detected by TripleTOF 6600 with an additional 2,544 specific proteins; and 87% of the proteins identified were common between the two ion mobility-equipped instruments (Exploris 480 and timsTOF Pro). In addition, Exploris and timsTOF pro detected ~13% specific proteins (692 and 638 proteins, respectively), suggesting the complementarity of different MS instruments for deep proteome coverage.

The Sankey plot (Fig. 4c) showed that proteins identified by newer generation instruments covered 90% of those identified by earlier generation instruments. Specifically, in the QE series, 96–100%, 94–96%, and 78–85% of proteins (sum of the red values in each row of the embedded table) in the high-intensity (> 66.67%), medium-intensity (33.33–66.7%), and low-intensity (< 33.33%) expression groups determined by the earlier generation instruments, respectively, were distributed in different intensity groups produced by the newer generation instruments. The vast majority of proteins in the three different intensity groups were reproducible in the corresponding groups with the same intensity level (Fig. 4c, values on the diagonal in the table) measured by newer generation instruments, especially those in the high-intensity group. Similar patterns were observed in the comparisons of the Fusion series, TOF-type instruments, and mass spectrometers with ion mobility. This assessment demonstrated that with continuous upgrading of instruments, the qualitative reproducibility of proteins with medium and high intensity is significantly increased (Supplementary Fig. 2a) and the detection of low-intensity proteins is improved, which would have a substantial effect on the reproducibility of proteome identification.

The median coefficient of variation (CV) of proteins in the global group of label-free quantitative proteomics generated by different instruments was approximately 30% (Fig. 4d and Supplementary Fig. 2b). Q Exactive, Q Exactive HF, and Exploris 480 showed a relatively low median CV (26%) in the global protein group, whereas Q Exactive HF-X showed a higher median CV (40%). Additionally, the median CV of proteins in the high-intensity group was lower than 20%, whereas proteins in the low-intensity group showed high variation, with the median CV value reaching up to 80%.

To objectively evaluate the performance of the MS platforms, we comprehensively considered the inherent differences of the Quartet standards and defined the SNR for quantitatively measuring the overall variations in the instrument or platform based on the inter- and intra-sample distances within principal component analysis dimensions (Fig. 4e, top panel). A higher value indicates that the inherent biological differences between samples have a greater contribution to the total variation relative to that caused by the state of the system or operation procedure. Thus, the SNR score can reflect the performance of an MS platform with respect to the reproducibility of the same sample and the capability of distinguishing different samples. The SNR scores of older versions of the instruments, such as Q-Exactive and TripleTOF 6600 that had lower protein identification coverage, were relatively higher (24 and 23.1, respectively) than those of newer generation versions such as the Fusion series, Q Exactive HF, and timsTOF (Fig. 4e, bottom panel). The proteins identified by earlier generations of mass spectrometers had relatively medium or high abundance, which increased the SNR. Although the newer generation spectrometers detected more proteins, especially low-abundance proteins, this resulted in more variation in biological quantification differences because of the increase in background complexity, thereby decreasing the SNR.

### Variation among laboratories

Quantitative evaluation based on the SNR, which ranged from 0.6 to 28, demonstrated substantial variation in the proteomic detection level of different platforms (Fig. 5a). An SNR of < 1 indicates that the variation introduced by system performance or operation procedure obscured inter-sample biological differences. Here, we judged results with an SNR of 0–2 as “ineligible,” 2– 10 as “average,” 10–20 as “good,” and > 20 as “excellent,” which accounted for 16.7%, 20.8%, 50%, and 12.5% of all 24 experiments, respectively. These results indicate that biological differences identified by most proteomics platforms (> 80%) were reliable and highlight the necessity of well-designed reference materials and suitable metrics in performance evaluation among different platforms.

**Fig. 5.**
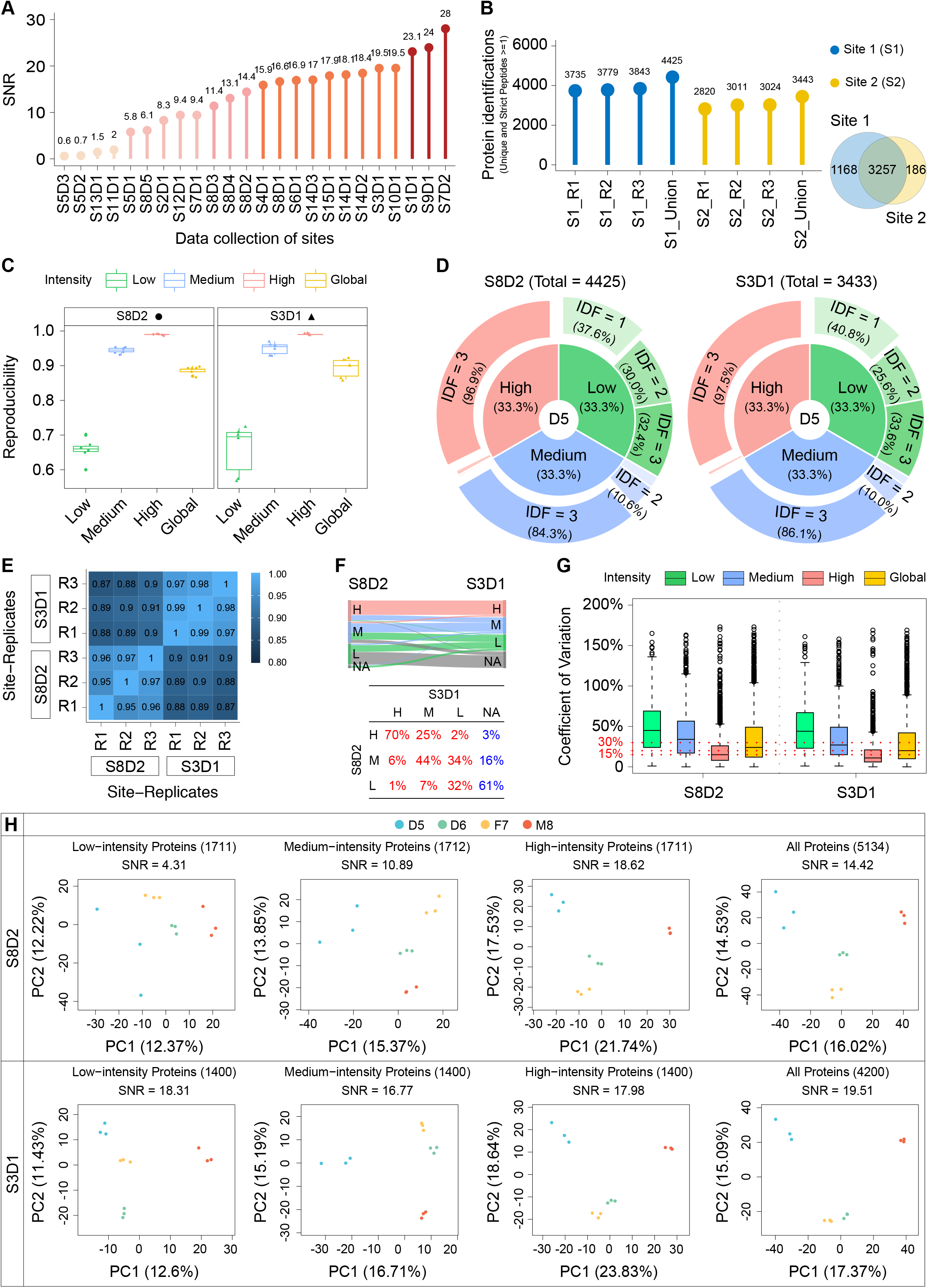
Variation among laboratories. **a**, SNR distribution of 24 datasets produced by the nine types of mass spectrometers in 15 laboratories. S: site; D: dataset. **b**, Number of proteins identified in S8D2 and S3D1 (left panel) and the Venn diagram of proteins identified from the two datasets (right panel). **c**, Reproducibility of detected proteins in low-intensity, medium-intensity, high-intensity, and global groups from the two datasets (left panel: S8D2; right panel: S3D1). **d**, Pie chart presenting the percentage of proteins with different identification frequency (IDF) in a specific experiment with three repeats; these proteins are divided into low-intensity, medium-intensity, and high-intensity groups (left panel: S8D2; right panel: S3D1). **e**, Pearson’s correlation coefficients matrix representing the correlations between different replicates from two laboratories. **f**, Sankey plot (top panel) depicting the flow direction of reproducibility for proteins within different (low, medium, high) intensity groups from S8D2 to S3D1. The corresponding and detailed percentages are summarized in the bottom table. **g**, CVs for proteins within different intensity groups and the global group of S8D2 and S3D1. **h**, Principal component analysis and SNR scoring for S8D2 and S3D1 using proteins within different intensity groups and the global group.

We next selected 2 of the 20 qualified experiments for qualitative and quantitative interlaboratory performance comparison. Two laboratories, Site 3 and Site 8, generated MS datasets S3D1 and S8D2, respectively, using Fusion Lumos. Taking sample D5 as an example, S8D2 included 4,425 proteins, which was 982 proteins more than those identified in S3D1 (Fig. 5b, left panel). Overall, 94.6% (3,257) of the 3,443 proteins in S3D1 were included in S8D2, with 1,168 and 186 site-specific proteins in S8D2 and S3D1 (Fig. 5b, right panel), respectively, demonstrating that the proteomics operation procedure at Site 8 was more suitable for in-depth proteome analysis. We then divided the proteins in S3D1 and S8D2 into low-, medium-, and high-intensity groups. Compared with S3D1, in-depth protein identification in S8D2 greatly improved protein reproducibility, especially in the low- and medium-intensity groups with a narrow interquartile range (Fig. 5c). The corresponding pie chart further strengthened this conclusion. For example, in the low-intensity group, 3.2% more proteins had an identification frequency > 1 in S8D2 than in S3D1 (62.4% vs. 59.2%; Fig. 5d). Pearson’s correlation coefficient between samples in S3D1 ranged from 0.97 to 0.99, which was superior to that of 0.95-0.97 in S8D2 (Fig. 5e). The Sankey plot indicated that proteins identified in S3D1 only covered ~73.6% (Fig. 5f, total proportion of the sum of the red values in each column of the table) of those identified in S8D2. Specifically, 97%, 84%, and 40% of proteins (Fig. 5f, the sum of the red values in each row of the table) in the high-, medium-, and low-intensity groups determined in S8D2 were distributed in different intensity groups than those in S3D1. These results indicate that compared with Site 3, Site 8 has advantages in detecting low-intensity proteins. The median CVs in the global group for label-free quantification in S3D1 and S8D2 were 20% and 24% (Fig. 5g and Supplementary Fig. 3a), respectively, indicating good quantification consistency.

Nevertheless, Site 3 showed a significant advantage over Site 8 in measuring the built-in difference of the Quartet standards with respect to the SNR (19.51 vs. 14.42) (Supplementary Fig. 3b); the SNRs gradually increased according to the increase in protein abundance (Fig. 5h). The SNR of the high-intensity sub-proteomes was comparable between S3D1 and S8D2, and although the SNR of the low- and medium-intensity sub-proteomes dramatically decreased to 4.31 and 10.89 in S8D2, it remained high for S3D1 (18.31 and 16.77, respectively). Therefore, we deduced that SNR scores were significantly more affected by the low- and medium-intensity sub-proteomes than by the high-intensity sub-proteome.

Collectively, these results demonstrate notable inter-laboratory differences in both qualitative and quantitative aspects, although most laboratories were able to identify biological differences; thus, it is necessary to perform inter-comparisons using the reference material. Such inter-laboratory comparisons can help laboratories validate the reliability of an SOP by determining the standard deviations of reproducibility and SNR and ultimately produce more confident results.

### Long-term stability of standards

To evaluate the stability of the Quartet standards, we conducted 15-month longitudinal monitoring for both peptides and proteins. All Quartet standards were stored at −80°C until analysis. Standards in protein form were tryptic-digested into peptides and then subjected to LC-MS/MS analysis, whereas standards in peptide form were directly loaded onto the LC-MS/MS system after dissolution. All stability evaluations were performed on the same Fusion Lumos instrument using the single-shot strategy. The protein and peptide form standards were tested monthly, generating a total of 120 MS files (2 types of standards × 4 samples × 15 months) (Fig. 6a).

**Fig. 6.**
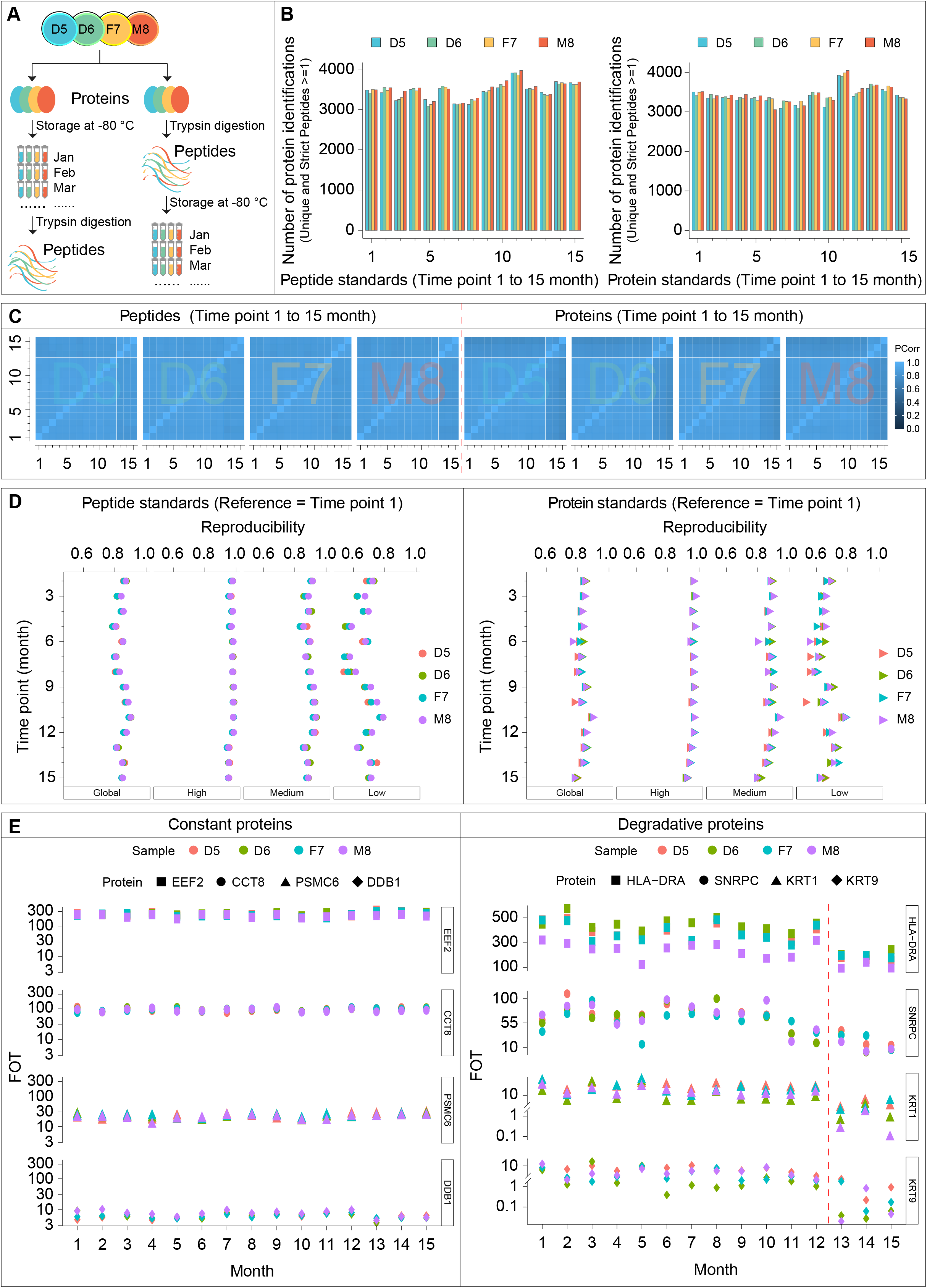
Long-term stability of standards. **a**, Longitudinal study design of stability evaluation of the Quartet standards. **b**, Number of proteins identified monthly at 15 different time points. Left panel: peptide standards; Right panel: protein standards. **c**, Pearson’s correlation coefficients matrix representing the correlation coefficients between the different mass spectrometry runs generated across 15 months using the Quartet standards (left panel: peptide standards; right panel: protein standards). **d**, Reproducibility of proteins within different intensity groups and the global group during 15-month monitoring (left panel: peptide standards; right panel: protein standards). **e**, Quantitative expression patterns of some proteins over 15 months. Left panel: proteins with high stability over 15 months; right panel: proteins with a degradation trend, especially after 12 months.

Little qualitative difference was found between the Quartet standards in peptide and protein forms with respect to the number of proteins identified at each monitoring point, with an average of 3,447 proteins identified at the 15 different time points (Fig. 6b). From a quantitative perspective, the proteomes had high correlations among experiments conducted in the first 12 months but were significantly weaker for the experiments performed in the last 3 months (Fig. 6c). This difference may have resulted from the degradation of peptides or proteins in the Quartet standards over time. Furthermore, we evaluated the reproducibility of proteins identified in the Quartet standards in peptide and protein forms, using the proteins identified in the first month as the reference (Fig. 6d). For standards in peptide form, the reproducibility of identified proteins showed high consistency, although there was a slight fluctuation trend, which was mainly caused by proteins in the low-intensity group. In the medium- and high-intensity protein groups, the inflection point of reproducibility occurred in the 13th month, which may have been due to the degradation of a few peptides. Comparatively, for standards in protein form, protein reproducibility started to show a relatively sharp decline by the 12th month for all three intensity groups. We speculated that this was also likely due to the degradation of more proteins. The stability of the prepared Quartet standards, in both peptide and protein forms, was high and similar for 1 year. Thereafter, the stability of standards in peptide form was better than in protein form. With respect to the quantitative expression patterns, some proteins in either medium- or high-intensity group, such as EEF2, CCT8, PSMC6, and DDB1, maintained high stability over 15 months, indicating their potential as internal standards (Fig. 6e, left panel). In addition, some proteins distributed in different intensity groups, such as HLA-DRA, SNRPC, KRT1, and KRT9, degraded over time, especially after 12 months (Fig. 6e, right panel). Taken together, these results demonstrate that both protein and peptide samples can be stably stored for 1 year at −80°C, with degradation occurring thereafter, and that the peptide form is more stable and reliable during storage than the protein form.

### Injection order contributes to proteome differences

We next evaluated the influence of the injection order on the quantitative capability of the MS systems. To this end, we compared the following three modes of injection order: random injective mode (RI), in which the Quartet standards were randomly injected into Q-Exactive HFX at any point within 1 week; continuous injection 1 (CI1), in which the Quartet standards were injected continuously into the same instrument (Q-Exactive HFX) in the order 5678-5678-5678; and continuous injection 2 (CI2), in which the Quartet standards were injected continuously into the same instrument (Q-Exactive HFX) in the order 555-666-777-888 (Fig. 7a). The proteins identified via the three modes showed more than 80% overlap, demonstrating that the injection order did not substantially affect the qualitative performance of the MS system (Fig. 7b). As shown in Fig. 7c, CI2 mode showed the highest quantification correlation among the three modes (Pearson’s correlation coefficient: 0.96-0.99), whereas the RI mode had the lowest quantitative performance (Pearson’s correlation coefficient: 0.90-0.98).

**Fig. 7.**
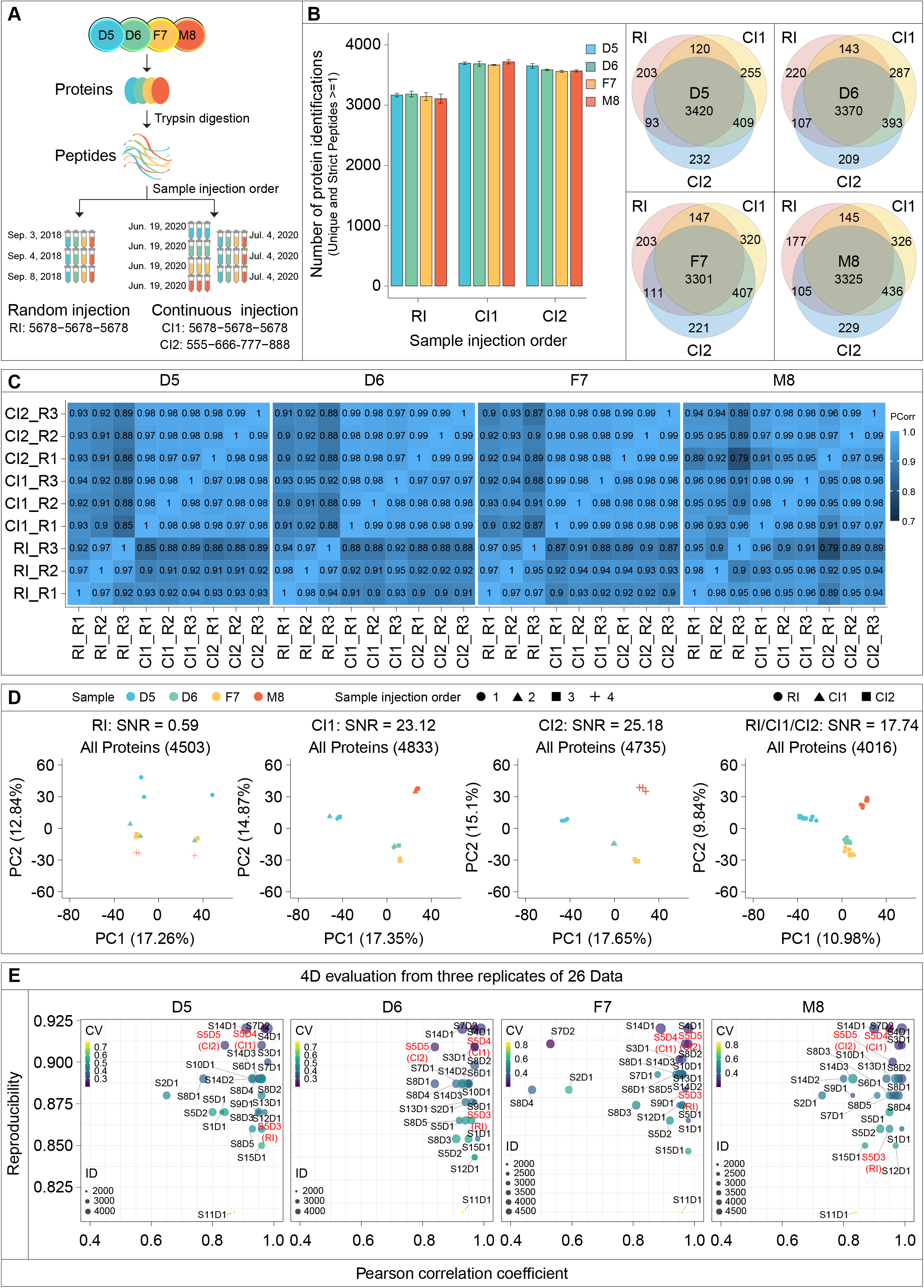
Injection order contributes to proteome differences. **a**, Experimental study design. **b**, Proteins identified in the Quartet samples using three different injection orders (left) and Venn diagram of the identified proteins (right). **c**, Pearson’s correlation coefficients matrix representing the correlation coefficients between different mass spectrometry runs generated by three different injection orders. **d**, Principal component analysis and signal-to-noise ratio (SNR) scores for datasets produced using different injection orders and the integrated dataset. **e**, Four-dimensional visualization map. RI: random injection order (5678-5678-5678); CI1: continuous injection order 1 (5678-5678-5678); CI2: continuous injection order 2 (555-666-777-888).

We also used the SNR indicator to evaluate the ability of the three injection modes to detect differences between the Quartet standard samples. The SNR of datasets produced in RI mode was only 0.59, indicating that the batch effect introduced by random injection had concealed biological differences. In contrast, the continuous injection strategy resulted in an excellent SNR score of 23.12 for CI1 mode and 25.18 for CI2 mode (Fig. 7d). Integration analysis of the data from the three injection modes showed an overall SNR of 17.74, suggesting that multiple repeats of samples reduce the batch effect. We thus proposed a four-dimensional visualization map (Fig. 7e) comprising protein identification, protein reproducibility, correlation among experiments, and experimental CV, which can be used to measure the proteomics operation level of each laboratory in a comprehensive manner, note the advantages and disadvantages of each laboratory, and highlight directions for further improvement.

### Absolute quantification of Quartet standards

To promote the large-scale, worldwide application of the Quartet as a proteome reference material, it is necessary to calculate the absolute number of the whole proteome (copy number per cell) according to the international metrological rules established by the International Organization for Standardization (ISO). To this end, we used the QconCAT method to measure the internal standard “anchor” proteins. We randomly selected 33 proteins with intensity-based absolute quantification (iBAQ) values distributed in four orders of magnitude as anchor proteins for absolute quantification and calibration of the copy numbers for the whole proteome. Representative unique peptides from these 33 anchor proteins were designed as QconCAT proteins by tandem fusion and split into 9 QconCAT proteins. Over 99% of the QconCAT proteins grown in the heavy isotope medium incorporated the labeled lysine, demonstrating the effectiveness of this approach.

To determine the absolute molar value of the C^13^-labeled QconCAT proteins, we synthesized the gold C^12^ GST peptide LLLEYLEEK (Institute of Metrology) and added gold peptides to nine tubes corresponding to the nine QconCAT proteins, which were digested with trypsin and loaded into the LC-MS/MS system for detection (Supplementary Fig. 4a). The molar amount of each QconCAT protein can then be calculated (Supplementary Table 4). For Quartet standard quantification, a dilution series of the C^13^-labeled QconCAT peptides was mixed with the Quartet peptide standard samples and then subjected to LC-MS/MS analysis (Fig. 8a). The corresponding heatmap portrays the linear trend of all identified C^13^-labeled anchor peptides (Fig. 8b and Supplementary Table 5). A dilution response curve of every C^13^-labeled anchor peptide (including GST peptide LLLEYLEEK) in the QconCAT proteins showed excellent linearity (Supplementary Fig. 4b, c), suggesting the good quantification accuracy of QconCAT proteins and the reliability of endogenous proteins in the Quartet anchored by QconCAT proteins.

**Fig. 8.**
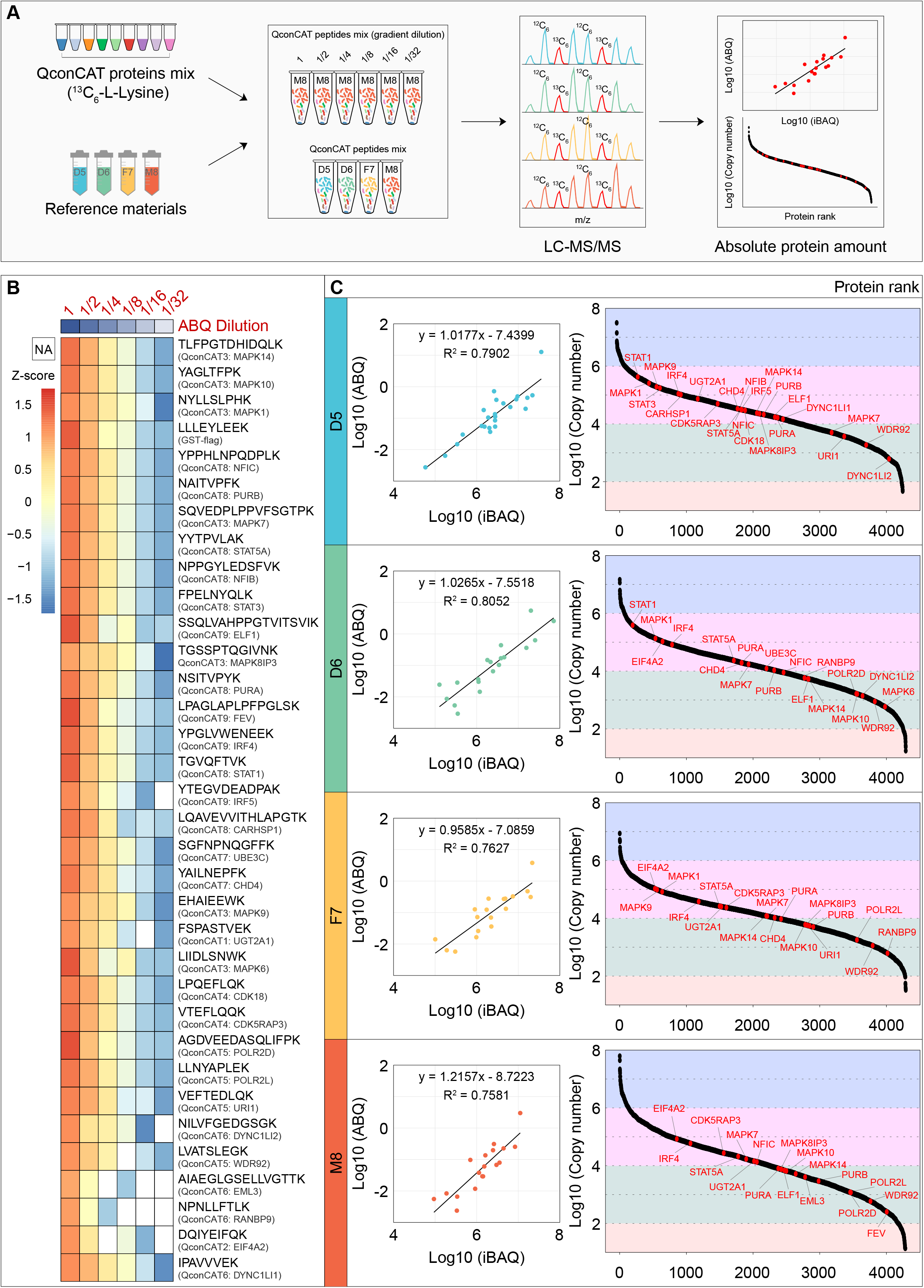
Absolute quantification of Quartet standards. **a**, Workflows for absolute quantification (ABQ) of the Quartet. **b**, Heatmap showing the linearity of peptide candidates as internal standards under a dilution series. **c**, Response curves between ABQs and iBAQs of the anchor proteins (internal standard) in each sample of the Quartet (left), and dynamic ranges of ABQs (copy number/cell) for each sample of the Quartet (right).

The absolute molar value of the corresponding proteins in the Quartet standard sample was determined based on quantification of the C^13^-labeled anchor peptides (Supplementary Table 6). The copy number of the 33 anchor proteins ranged from 10^2^ to 10^6^ copies/cell. We aligned the absolute copy numbers (ABQ) of the anchor proteins to their corresponding iBAQ values and found that the ABQs and iBAQs were well-correlated (R^2^ = 0.7581-0.8052) (Fig. 8c), demonstrating consistency between relative quantification (iBAQ) and absolute quantification (copy number). Finally, we quantified the abundance of more than 4,000 proteins in each sample of the Quartet reference materials by aligning the proteome to the anchor proteins (Supplementary Table 6). Their dynamic ranges spanned over ~7 orders of magnitude from 10^1^ to 10^8^ copies/cell. Thus, absolute quantification of the Quartet proteome can provide reference datasets for QC and quality assessment of an LC-MS/MS platform for both basic research and clinical applications.

## Discussion

Quantitative proteomics is playing an increasingly prominent role in clinical and basic research and has been gradually introduced into clinical practice. Proteomic molecular subtyping research led by the Clinical Proteomic Tumor Analysis Consortium (CPTAC) and the Chinese Human Proteome Project (CNHPP) consortia dissected the proteomes of several cancer types, including brain cancer^33^, hepatocellular carcinoma^34, 35^, lung cancer^36^, lung adenocarcinoma^37, 38^, gastric cancer^39^, colon cancer^40–42^, clear cell renal cell carcinoma^43^, endometrial carcinoma^44^, ovarian cancer^45^, and breast cancer^46, 47^. These projects proposed druggable targets and indicators for selecting clinical treatment strategies. Moreover, proteomics analyses of blood, urine, and cerebrospinal fluid samples have facilitated the discovery of several molecular markers of human diseases^48–50^. However, to guarantee the reproducibility of the molecular subtyping and markers revealed by proteomics, it is vital to conduct a thorough and objective multi-platform assessment of the utility of particular proteomics technologies.

As early as 2004, the MAQC project led by the FDA took the first step toward addressing the reproducibility crisis in genomics. The project developed RNA reference materials by combining biologically different RNA sources and known titration differences for cross-platform assessment, generating a rich dataset and revealing promising results regarding the consistency of microarray data between laboratories and across platforms^17, 18^. Despite similar concerns regarding the accuracy of published proteomics data within the literature, along with a lack of universal standard materials, SOPs, and benchmark datasets, reproducibility of proteome datasets and the reliability of the associated biological insights obtained from these data remain significant issues to be addressed.

The Proteomics Standards Research Group (sPRG) generated a well-defined protein standard with a mixture of 49 human proteins for use in proteomics (http://www.abrf.org/sPRG). Among 74 participating laboratories, 66% identified at least 30 out of the 49 proteins, with only 8% identifying < 10 of the proteins^51^. Among the slightly different analysis/identification methods used, no single approach was identified as superior. In addition, using the sPRG study results as a foundation, Sigma-Aldrich released the first commercially available proteomics standard in 2006, the Universal Proteomics Standard (UPS1). To account for the broad dynamic range found in most proteomics samples, Sigma-Aldrich subsequently introduced the Proteomics Dynamic Range Standard (UPS2) using the same proteins. The development of USP1/2 enabled evaluation of LC-MS/MS system performance across different laboratories and investigation of the sources of variation^22, 52^. However, USP1/2 cannot truly simulate the entirety of the human proteome, making these standards unsuitable for the QC experiments required for large-scale clinical proteomics research^53^.

Here, we performed a comprehensive multi-platform assessment, including protein detection and quantification, using label-free LC-MS/MS-based proteomic profiling involving nine types of major mass spectrometers and 15 proteomics laboratories, focusing on inter-instrument/laboratory variability and the effects of injection order on the discovery of biological differences. Our benchmark study produced well-characterized reference materials, referred to as “the Quartet,” which contained differences between samples as “built-in truths” at both the peptide and protein levels for evaluating the ability to detect differential proteomes. Only 6% of the proteins were steadily identified in all MS measurements. In contrast, more than 50% of the proteins were detected in less than 10% of the MS measurements. Consistent results were observed for all four standard samples in the Quartet derived from a Chinese family. These percentage objectively reflected the random nature of LC-MS/MS-based proteomics and indicated the bias level between different technical MS repeats^24, 25, 52^. The percentage of reproducible DEPs identified was dramatically decreased by two magnitudes compared with that of protein identification. For example, 0.03-0.3% of DEPs were consistently discovered in all surveyed MS measurements and 6-9% of the DEPs were discovered in 50% of the surveyed MS measurements. Accordingly, we recommend performing at least three technical repeats for reliable identification and many more repeats for reliable quantification.

Another advantage of the Quartet standard is the ability to identify intrinsic proteomic differences among the four samples tested herein, which facilitated evaluation of the performances of MS-based identification/quantification in distinguishing biological samples^54^. We introduced an SNR score algorithm to objectively evaluate the performance of the MS platforms. Surprisingly, the different platforms from the 15 laboratories markedly varied in terms of SNR score, and the worst dataset was not able to clearly separate the inter- and intra-sample differences of the Quartet. Although this result may not be representative, it may partially reflect the current situation of proteomics research, further demonstrating the necessity of applying a reference material for performance evaluation of different platforms. Further analysis indicated that the low- and medium-intensity proteome regions, but not the high-intensity proteome region, were largely responsible for the final SNR score, even though the high-intensity proteins accounted for the majority of the proteome abundance.

The stability of protein samples and their proper storage have remained unresolved issues in the field of proteomics^55^. We found good stability of both protein and peptide samples for at least 1 year with storage at −80°C and demonstrated that it is better to store the sample in the peptide form. Furthermore, this Quartet standard enabled assessment of sample stability at different facilities under different storage conditions and optimization of the storage approach accordingly. The longitudinal monitoring results also highlighted the importance of thoroughly assessing the proteome of the standard and annually updating the reference datasets.

This study also draws attention to the impact of the injection order on the quantification performance of LC-MS/MS. Even though batch effects are common concerns for all omics technologies, we were still surprised to find that the random injection approach, in which target injections were separated by other irrelevant samples, significantly decreases the capability of distinguishing inter- and intra-sample differences. Therefore, we highly recommend that samples for a given project be continuously injected without interruption. If interruption is inevitable, more biological and technical replicates should be performed to overcome this limitation.

In summary, we established a proteome standard reference materials, termed the Quartet, with absolute protein expression reference values. A total of 816 MS files were thoroughly profiled, and the qualitative and quantitative characteristics of the Quartet standard samples were determined, providing valuable reference datasets for proteomics analyses. As the Quartet standards were developed within the MAQC-IV project, the DNA, RNA, and metabolite forms from the same batch of samples are also available, thereby allowing for trans-omics integration of the corresponding genome, transcriptome, and metabolome datasets. We believe that the Quartet protein standards, along with other biomaterial forms, will have broad applications in basic research and translational medicine.

## Online Methods

### Sample collection and preparation

Lymphoblastoid cells provided by a family from the FDU Taizhou cohort, which consisted of a father (F7), mother (M8), and a pair of twin daughters (D5 and D6), were used for the establishment of immortalized cell lines^29–31^. All cells were cultivated in RPMI-1640 medium (HyClone, Logan, UT) supplemented with 10% fetal bovine serum (HyClone) and incubated at 37°C in 5% CO_2_. Protein extraction from the cells and peptide digestion were uniformly performed in the National Center for Protein Sciences (NCPSB) laboratory (Beijing, China). Two types of prepared reference materials derived from 2 × 10^9^ cells of each of the four family members were packed into 1,000 EP tubes with specific color labels (D5: blue, D6: green, F7: yellow, and M8: red) and stored in a −80°C freezer. For generating the protein standard material, aliquots of 100 μg total protein were stored in each tube. For generating the peptide standard material, aliquots of 10 μg total tryptic peptide were stored in each tube. We distributed these aliquots to 15 laboratories throughout six cities in China (Fig. 1a).

### Preparation of QconCAT proteins

QconCAT proteins were prepared according to the procedures reported by Ding et al.^56^. Briefly, the QconCAT cDNA was reverse-translated from amino acid sequences of the selected QconCAT tryptic peptides and chemically synthesized (Generay Biotech, Shanghai, China) before cloning into the pGEX4T-1 vector (Addgene, Watertown, MA). The GST-QconCAT plasmids were then transformed into *Escherichia coli* BL21 cells (TransGen Biotech, Beijing, China) for protein expression. A fresh *E. coli* BL21 culture was inoculated into 5 mL of heavy SILAC Dulbecco’s modified Eagle medium (^13^C6 lysine, without glutamine; Invitrogen, Carlsbad, CA) and grown at 37°C for 16 h. Isopropyl-β-D-1-thiogalactopyranoside (0.4 mM; Sigma-Aldrich, St. Louis, MO) was added to the bacterial culture when the absorbance at 600 nm reached ~0.5 to induce QconCAT protein expression at 37°C for 3 h. The expression and identity of the recombinant protein was verified by mass spectrometry. The BL21 cells were collected, suspended in NETN buffer (150 mM NaCl, 1 mM EDTA, 50 mM Tris-HCl, 1% NP-40, with protease inhibitors), and lysed on ice by sonication. The lysate was then centrifuged at 60,000 rpm for 10 min, and the supernatant was collected to purify GST-QconCAT recombinant protein using GSH beads. The purified GST-QconCAT proteins were eluted by elution buffer (10 mM glutathione, 50 mM Tris-HCl, 5% glycerol) and stored at −80°C until use.

### Protein extraction and tryptic digestion

The cells were lysed in lysis buffer (8 M urea, 100 mM ammonium bicarbonate, pH 8.0) supplemented with protease inhibitors for 10 min on ice, and then sonicated for 3 min (2 s on and 2 s off) on ice and centrifuged at 16,000 × *g* for 20 min to remove the cell debris. The supernatants were collected, and the protein concentration was measured using a bicinchoninic acid protein assay. Extracted proteins were loaded into 10-kD Microcon filtration devices (Millipore, Burlington, MA), centrifuged at 16,000 × *g* for 20 min, and washed twice with urea lysis buffer and twice with 50 mM NH_4_HCO_3_. The samples were then digested using trypsin at an enzyme:protein mass ratio of 1:25 overnight at 37°C, after which peptides were extracted and dried (SpeedVac; Eppendorf, Hamburg, Germany)^57^. Two types of proteome standards in peptide and protein forms were uniformly prepared and sub-packed into EP tubes, followed by storage in a −80°C freezer. Before use, the protein standards were digested into peptides.

### One-dimensional reversed-phase separation

The dried peptides were loaded into a home-made Durashell reverse-phase column (2 mg packing, 3 μm, 150 Å, in a 200 μL tip; Agela, Torrance, CA), and then sequentially eluted with different gradients of elution buffer containing mobile phase A [2% acetonitrile (ACN), adjusted to pH 10.0 using NH_3_·H_2_O) and different concentrations of mobile phase B (98% ACN, adjusted to pH 10.0 using NH_3_·H_2_O). The different fractions (6, 12, 18, or 30) were then vacuum dried for sub-sequential MS analysis.

### LC-MS/MS analysis

Peptide samples were analyzed on Orbitrap Fusion Tribrid series (Fusion and Fusion Lumos), Q Exactive hybrid quadrupole-Orbitrap series (Q-Exactive, Q-Exactive Plus, Q-Exactive HF, Q-Exactive HF-X, and Exploris 480 with ion mobility) (both from Thermo Fisher Scientific, Waltham, MA), Sciex Triple-TOF 6600 System (SCIEX, Foster City, CA), or Bruker timsTOF Pro with ion mobility (Bruker Daltonics, Bremen, Germany) mass spectrometers, coupled with an Easy-nLC 1000 nanoflow LC system (Thermo Fisher Scientific), Ultra Plus NanoLC 2D HPLC system (Eksigent Technologies, Dublin, CA), or Elute-HT UHPLC system (Bruker Daltonics). Dried peptide samples redissolved in Solvent A (0.1% formic acid in water) were loaded onto a 2 cm self-packed trap column (ReproSil-Pur C18-AQ; 100 μm inner diameter, 3 μm; Dr. Maisch, Ammerbuch, Germany) using Solvent A and separated on a 150 μm inner diameter column with a length of 15 cm (ReproSil-Pur C18-AQ, 1.9 μm; Dr Maisch) using a gradient (buffer A: 0.1% formic acid in water; buffer B: 0.1% formic acid in 80% ACN) at a constant flow rate of 600 nL/min. MS was operated under data-dependent acquisition mode. All parameters were set according to the requirements of the respective instruments.

### MS database search

MS raw files generated by LC-MS/MS were searched against the human National Center for Biotechnology Information Refseq protein database (updated on April 7, 2013; 32,015 entries) using Firmiana 1.0 enabled with Mascot 2.3 (Matrix Science Inc., London, UK). The protease was trypsin/P, and up to two missed cleavages were allowed. Carbamidomethyl was considered a fixed modification. For the proteome profiling data, variable modifications were oxidation and acetylation (Protein N-term). Proteins with at least one unique peptide with a 1% false-discovery rate at the peptide level and a Mascot ion score greater than 20 were selected for further analysis.

### Peptide and protein quantification

Peptide quantification was performed according to the peak area. Proteins were quantified using the label-free iBAQ approach. The fraction-of-total (FOT) is used to represent the normalized abundance of a particular protein, which was defined as a protein’s iBAQ value divided by the total iBAQ of all identified proteins within one sample. The FOT was multiplied by 105 for ease of presentation in figures.

### Differential protein analysis

The fold change in expression level was used to determine whether proteins were differentially expressed between samples. Proteins with expression level changes greater than 2-fold were considered to be upregulated or downregulated. The proteins identified from MS runs at the participating laboratory for each sample in the Quartet were divided into multiple groups according to the confidence intervals of their occurrence frequency in all experiments (observation times/total measurement times).

### Statistical analysis

Pearson’s correlation coefficient (r) was calculated using the cor.test function in the R software. The protein overlap rate was used as a measure of reproducibility, and the CV was used as a measure of quantification consistency. For the heatmap, each protein expression value in the global proteomic expression matrix was transformed into a Z-score across all samples. For protein-wise clustering, the distance was set as “Euclidean” distance and the weight method was set to “complete.” The Z-score-transformed matrix was clustered using the R package pheatmap (version 1.0.12). The SNR was calculated from the inter- and intra-sample distances within principal component analysis dimensions (Fig. 4e, upper panel). All other statistical analyses were performed with RGUI version 3.6.

## Data and code availability

All proteomic raw data have been uploaded to the iProx Consortium (https://www.iprox.org/) with the subproject ID (IPX0002859000). In addition, all proteomics raw data have been deposited at the Firmiana platform (a one-stop proteomic cloud platform; https://phenomics.fudan.edu.cn), and the qualified profiling datasets were processed at the platform.

## Acknowledgements

This work was supported by the National Key R&D Program of China (2017YFA0505102, 2016YFA0502500, 2018YFA0507501, 2017YFC0908404, 2020YFE0201600, 2018YFE0201603, 2017YFA0505101), the National Natural Science Foundation of China (31770886, 31972933, 31700682), the Shanghai Municipal Science and Technology Major Project (2017SHZDZX01), the Major Project of Special Development Funds of Zhangjiang National Independent Innovation Demonstration Zone (ZJ2019-ZD-004), and the Fudan Original Research Personalized Support Project.

## Author contributions

Conceptualization, C.D., J.Q, F.H, Y.Z., and Proteomic MAQC Consortium; Methodology, S.T., D.Z., Y.Y., M.L., Y.W., L.S., Z.Q., X.L., S.J., Yan Li, L.L., S.W., Proteomic MAQC Consortium, Y.Z., F.H, J.Q., and C.D.; Software, S.T., D.Z., Y. Liu, Yao Li, J.Q., and C.D.; Formal Analysis, S.T., D.Z., Y.Y., Y.Z., F.H., J.Q., and C.D.; Investigation, S.T., D.Z., Y.Y., M.L., Y.W., L.S., Z.Q., X.L., Y. Liu, Yao Li, S.J., Yan Li, L.L., S.W., Y.Z., and C.D.; Resources, S.T., D.Z., Y.Y., Proteomic MAQC Consortium, Y.Z., F.H., J.Q., and C.D.; Data Curation, S.T., D.Z., Y.Y., Proteomic MAQC Consortium, Y.Z., F.H., J.Q., and C.D.; Writing–Original Draft, S.T., D.Z., Y.Y., and C.D.; Writing– Review & Editing, S.T., Y.Z., F.H., J.Q., and C.D.; Visualization, S.T., D.Z., Y. Liu, Yao Li, Yan Li, L.L., and C.D.; Project Administration, S.T., D.Z., Y.Z., F.H., J.Q., and C.D.; Funding Acquisition, Y.Z., F.H., J.Q., and C.D.

## AUTHORS

**Project leader:** Chen Ding

**Manuscript preparation team leader:** Chen Ding

**Scientific management:** Chen Ding

**Proteomic Massive Analysis and Quality Control Consortium:**

Chen Ding^1^, Jun Qin^2^, Fuchu He^1,2^, Yuanting Zheng^1^, Sha Tian^1^, Dongdong Zhan^2,12^, Ying Yu^1^, Yunzhi Wang^1^, Zhaoy Qin^1^, Yang Liu^1^, Yao Li^1^, Yan Li^1^, Lingling Li^1^, Qingyu He^3^, Xingfeng Yin^3^, Lunzhi Dai^4^, Haiteng Deng^5^, Chao Peng^6^, Ping Wu^6^, Minjia Tan^7^, Jing Jiang^8^, Yaoyang Zhang^9^, Yunxia Li^9^, Wenqin Liu^10^, Wei Chen^11^, Rui Wang^11^, Jin Zi^13^, Qidan Li^13^, Mingzhou Bai^13^, Zeng Wang^13^, Zhanlong Mei^13^, Zhongyi Cheng^14^, Jun Zhu^14^, Xuemei Wu^15^, Xing Yang^16^, Yue Zhou^17^

### Affiliations

^1^Fudan University, Shanghai 200433, China.

^2^National Center for Protein Sciences (NCPSB), Beijing 102206, China.

^3^Jinan University, Guangzhou 510632, China

^4^Sichuan University, Chengdu 610017, China

^5^Tsinghua University, Beijing 10084, China

^6^National Center for Protein Science Shanghai, Shanghai 200237, China

^7^Shanghai Institute of Materia Medica Chinese Academy of Sciences, Shanghai 201203, China

^8^The First Affiliated Hospital of Medical School of Zhejiang University, Zhejiang 310000, China

^9^Interdisciplinary Research Center of Biology and Chemistry, Shanghai 201210, China

^10^Asia Pacific Application Support Center, AB SCIEX, Shanghai 201101, China

^11^APTBIO, shanghai 200233, China

^12^Beijing Guhai Tianmu Biomedical Technology Co., Ltd., Beijing 100093, China.

^13^BGI-Shenzhen, Shenzhen, Guangdong 518083, China

^14^PTM Bio, Hangzhou 310018, China Shanghai Office, Bruker Daltonics Inc., Shanghai 200233, China

^15^Shanghai Office, Bruker Daltonics Inc., Shanghai 200233, China

^16^Shanghai iProteome Biotechnology Co., Ltd, Shanghai 201203, China

^17^Thermo Fisher Scientific(China) Co. Ltd, Shanghai 2100120, China

