## Supplementary figure for "Quartet protein reference materials and datasets for multi-platform assessment of label-free proteomics": Supplementary information.pdf

### Supplementary figure 1

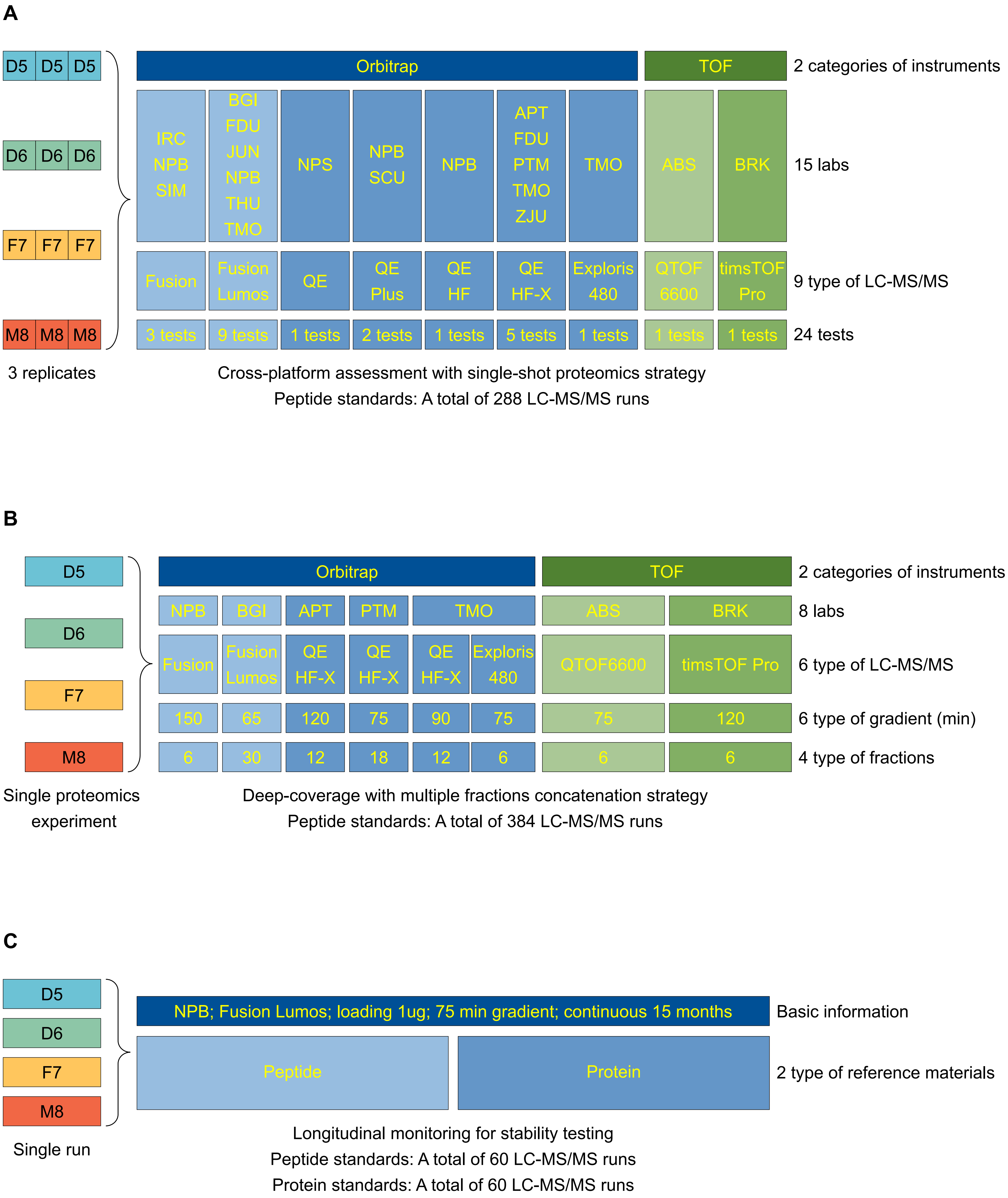

### Supplementary figure 2

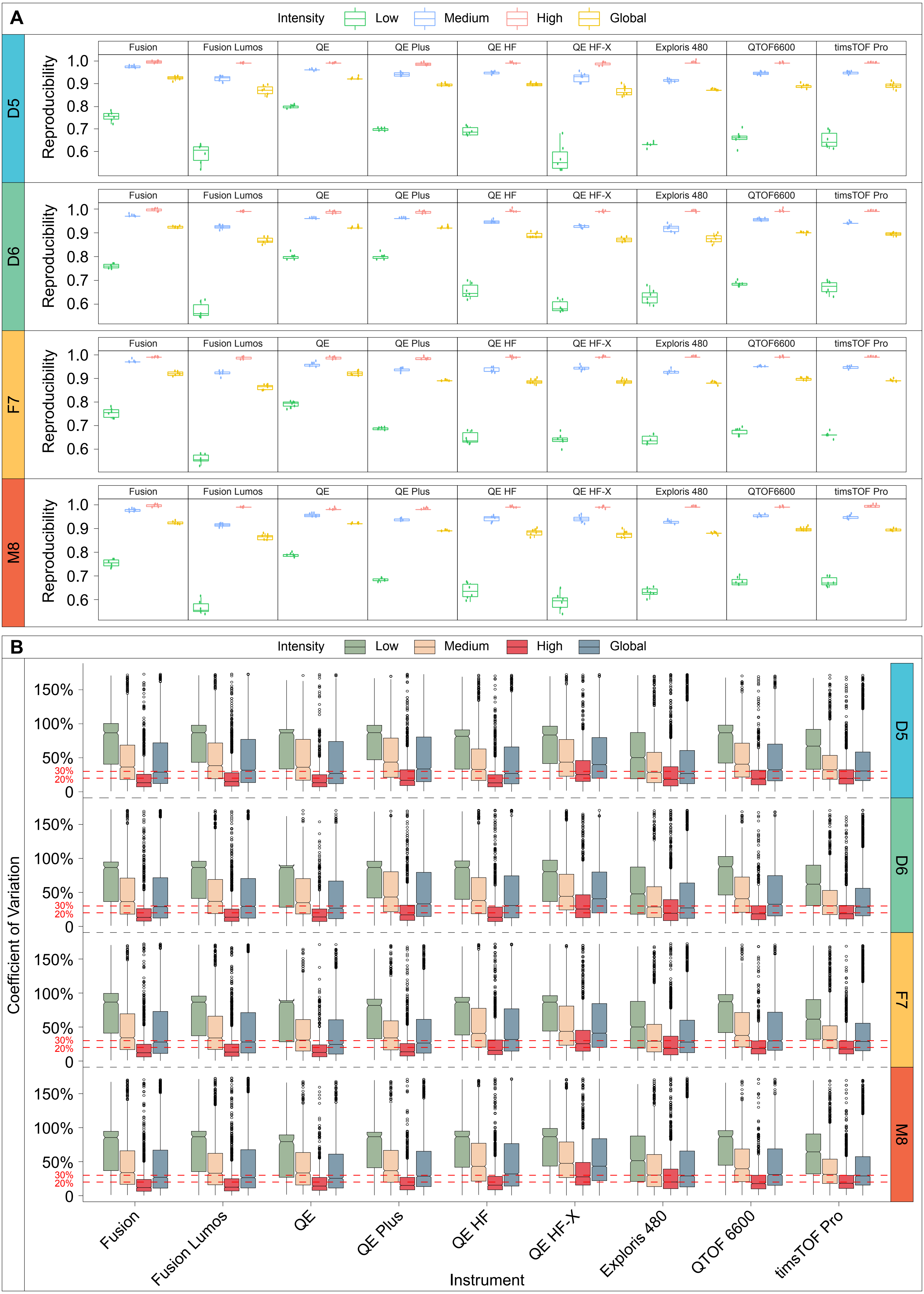

### Supplementary figure 3

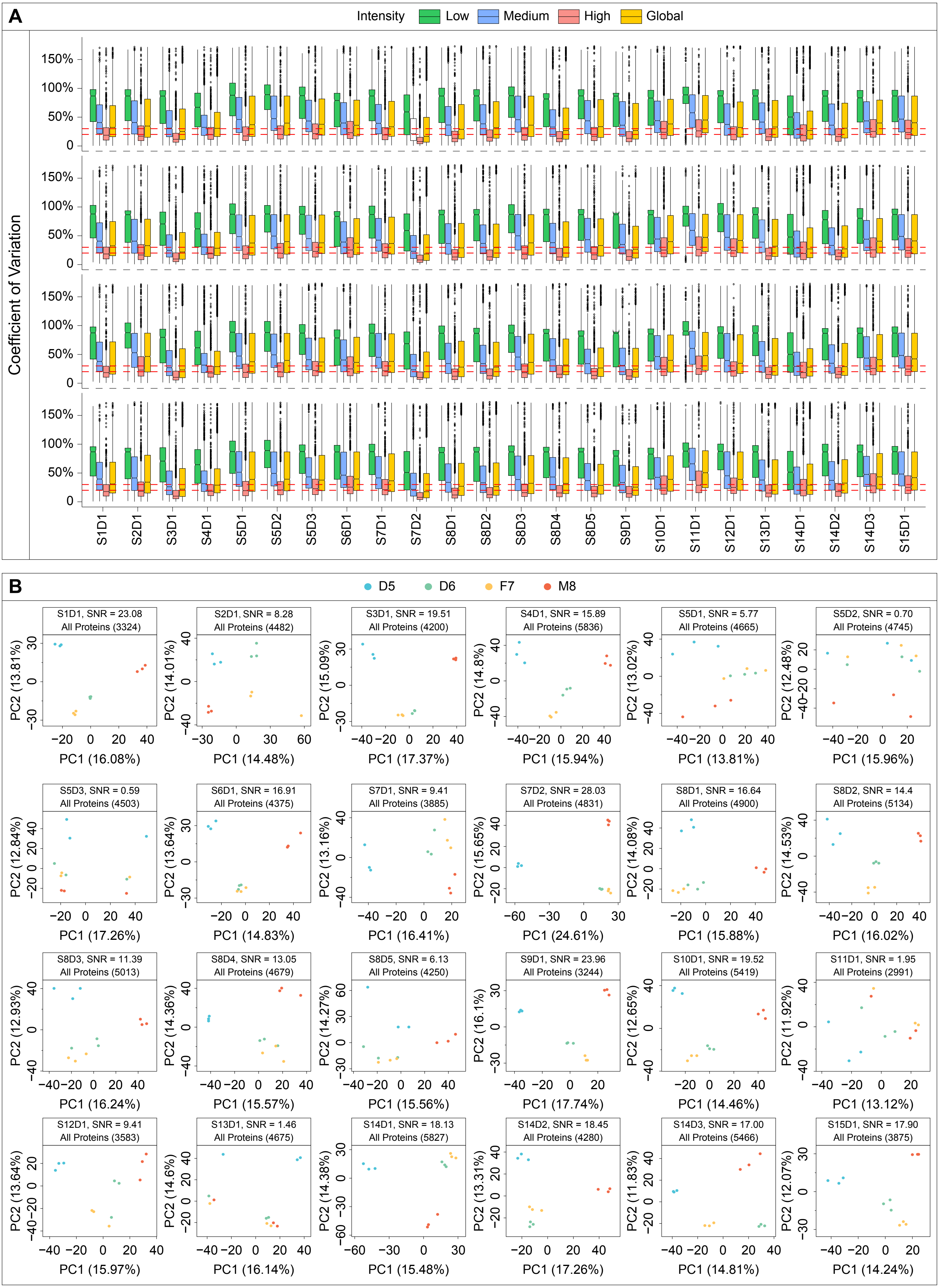

### Supplementary figure 4

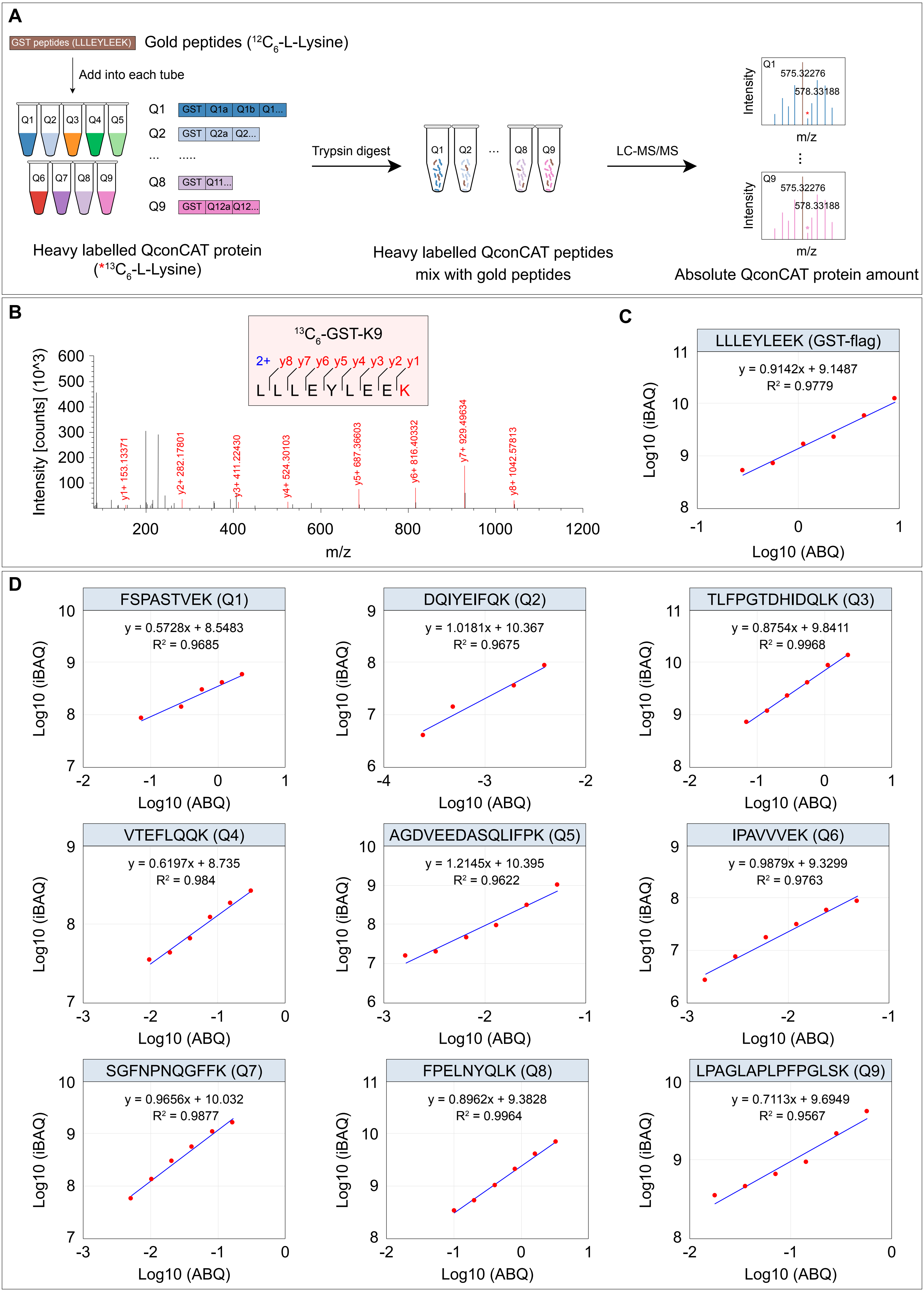

**Supplemental Information legends**

**Supplementary Fig. 1 | MS files (N = 792) used to generate standard reference material datasets, Related to Fig. 1. a**, Cross-platform assessment with the single-shot proteomics strategy (peptide standards with a total of 288 LC-MS/MS runs). **b**, Deep coverage with a multiple fraction concatenation strategy (peptide standards with a total of 384 LC-MS/MS runs). **c**, Longitudinal monitoring for stability testing (peptide standards: total of 60 LC-MS/MS runs; protein standards: total of 60 LC-MS/MS runs)

**Supplementary Fig. 2 | Quantitative reproducibility and variation analysis among nine conventional instruments, Related to Fig. 4. a**, Reproducibility of detected proteins from a quantitative perspective (low-intensity, medium-intensity, high-intensity, and global groups) analyzed by nine conventional instruments in each sample of the Quartet. **b**, Quantitative variation of each sample of the Quartet in the low-intensity, medium-intensity, high-intensity, and global groups of nine conventional instruments (to facilitate overall comparison, the CVs of D5 among nine conventional instruments are repeated here).

**Supplementary Fig. 3 | Quantitative evaluation based on CVs and SNR, Related to Figure 5. a**, Quantitative variation of each sample of the Quartet in the low-intensity, medium-intensity, high-intensity, and global groups of 24 datasets produced by nine types of mass spectrometers across 15 laboratories (to facilitate overall comparison, the CVs of D5 among S8D2 and S3D1 are repeated here). **b**, Principal component analysis and SNR scoring of all proteins groups from 24 datasets produced by nine types of mass spectrometers across 15 laboratories (to facilitate overall comparison, the principal component analysis and SNR scoring in S8D2 and S3D1 are also repeated here).

**Supplementary Fig. 4 | Absolute quantification of the QconCAT proteins. a**, Workflows for absolute quantification (ABQ) of the QconCAT proteins. **b**, MS/MS spectra (left panel) and dilution response curve (right panel) of the $C^{13}$-labeled gold peptide (LLLEYLEEK) in GST-flag of the QconCAT proteins. **c**, Dilution response curve of the $C^{13}$-labeled representative anchor peptide in every QconCAT protein.
